# Implicit bias is strongest when assessing top candidates

**DOI:** 10.1101/859298

**Authors:** Emma R Andersson, Carolina Hagberg, Sara Hägg

**Affiliations:** Department of Biosciences and Nutrition, Karolinska Institutet, Sweden; Department of Cell and Molecular Biology, Karolinska Institutet, Sweden; Department of Medicine, Solna, Karolinska Institutet, Sweden; Department of Medical Epidemiology and Biostatistics, Karolinska Institutet, Sweden

**Keywords:** equality, diversity, life science, peer review, bibliometry, faculty positions, multivariable analysis

## Abstract

**Background:** Academic life is highly competitive and expectations of fair competition underlie the assumption that academia is a meritocracy. However, implicit bias reinforces gender inequality in all peer review processes, unfairly eliminating outstanding individuals and depleting academia of diversity. Here, we ask whether applicant gender biases reviewer assessments of merit in Sweden, a country that is top ranked for gender equality.

**Methods:** We analyzed the peer review procedure for positions awarded at a Swedish medical University, Karolinska Institutet (KI), during four consecutive years (2014-2017) for Assistant Professor (n=207) and Senior Researcher (n=153). We derived a composite bibliometric score to compute productivity, and compared this to subjective external (non-KI) peer reviewer scores on applicants’ merits to test their association for men and women, separately.

**Results:** Men and women with equal merits are not scored equally by reviewers. Men generally have stronger associations (steeper slopes) between computed productivity and subjective external scores, meaning that peer reviewers suitably “reward” men’s productivity with increased merit scores. However, for each additional composite bibliometric score point, women applying for Assistant Professor positions only receive 58% (79% for Senior Researcher) of the external reviewer score that men received, confirming that implicit bias affects external reviewers’ assessments. As productivity increases, the difference in merit scores between men and women increases.

**Conclusions:** Accumulating bias impacts most strongly in the highest tier of competition, the pool from which successful candidates are ultimately chosen. Gender bias is apparent in external peer review processes of applications for academic positions in Sweden, and is likely to reinforce the unbalanced numbers of professorships in Sweden.

## INTRODUCTION

Fostering groundbreaking research requires identification of the best ideas and individuals. Competition in academia is fierce, and only 47% of those that are awarded a PhD pursue a career within science after their PhD, and only 0.45% become professor ^1^. Resources to support researchers are limited, and thus it is essential that the best candidates be identified, in order to use resources wisely. However, implicit bias may limit career progression for women or minorities. Recent data suggest both overt and implicit bias is generally decreasing ^2^, yet women still make up only one third of all professors in European countries ^3–5^.

Implicit bias affects multiple aspects of a scientific career, including decision making in recruitment ^6,7^, grant awarding ^8–10^, and citations ^11^. An entry-level group leader position in academia is typically awarded following peer review of the applicants and their proposed projects, assessing quality, innovativeness, feasibility and potential. In order to assess whether, and how, bias impacts upon career progression, we have analyzed the peer-review procedure for positions awarded at a Swedish medical University, Karolinska Institutet (KI), over a period of 4 years, including 1187 eligible applicants, of which the top 30 % (360 applicants) were selected for a Step 2 peer review by external non-KI reviewers in Sweden. By computing a composite bibliometric score, which includes seven publication parameters, as a proxy for productivity, we quantify the applicants’ records of accomplishment. Composite bibliometry is a powerful tool to assess multiple bibliometric parameters that has previously been used to objectively measure productivity ^8,10^, and is a strong predictor of scientific quality, for example more accurately identifying Nobel Prize winners than citations alone ^12^. The composite bibliometric score was compared to the external reviewer scores assigned during Step 2 of the peer-review process, to assess whether men and women of equal productivity were considered equally merited. Our analysis shows that men are generally awarded higher merit scores by reviewers for equal productivity (assessed by the composite score), and we show that this discrepancy in scoring is greatest at the top of the competition. While much has improved with regards to equal opportunities, continued efforts to eliminate implicit bias are clearly needed to achieve parity, in particular in assessing top candidates.

## METHODS

### Data

Each year since 2014, KI has announced position grants to recruit Assistant Professors and Senior Researchers within a Career Ladder scheme (https://ki.se/en/about/faculty-funded-career-positions). Assistant Professor in the KI Career Track 2014-2018 was a 4 year position as independent group leader/PI, typically the first faculty appointment. Senior Researcher was a 5 year position, typically awarded after completion of an Assistant Professor position, the equivalent of Associate Professor. This grant scheme entails a three-step process, in which applicants submit a CV and a project plan. In Step 1, the top 30 % of applicants are selected for external peer review. In Step 2, these top applications are assessed by external reviewers. Six external reviewers, professors at other Swedish universities, with equal gender distribution, score each application according to instructions provided by KI (Appendix). They provide one score for merits (based on the CV) and one for the project plan, returning a ranking to KI. In Step 3, the top 20 or so applicants are interviewed by KI professors, competing for one of 7-11 positions for Assistant Professor, or one of 6-8 positions as Senior Researcher. There is no selection based on research field or type of research.

The data in this study includes information on applications for KI Career Ladder positions between 2014-2017; total number of eligible applicants, number sent to external review, number that went to interview, and number awarded. All applications that were sent to external review were included in the following analysis. In total, 360 applications from Step 2 (external review) were assessed, of which 207 were applications for Assistant Professor and 153 for Senior Researcher. The data included gender and the average external non-KI reviewer scores of the applicant’s merits. Data were provided by KI registrar.

The 2014 data for Assistant Professors have been analysed and reported once before using a slightly different composite bibliometric score ^8^.

### Composite bibliometric score

The composite bibliometric score was derived by the KI library analytic team based on articles and reviews published from 1995 to the time of application and available in the Web of Science. The score consists of seven parts: 1) the number of publications, 2) the total number of citations to those publications, 3) the share of publications where the applicant was the first author, 4) the share of publications where the applicant was the last author, 5) the H index, 6) the share of publications in high impact journals within its field, and 7) a binary indicator for having any publication in a high impact journal overall. Within a field, the top two journals by Journal Impact Factor were considered high impact and overall the top 30 journals were considered high impact. Citations were retrieved from the Web of Science and Journal Impact Factors from Journal Citation Reports, both maintained by Clarivate Analytics. Each part of the score was log-transformed and then standardized so that the smallest value corresponds to a standardized value of 0 and the largest value to a standardized value of 1. The composite score is the sum of these seven standardized variables.

### Statistical analysis

Linear regression models were used to quantify the association between the composite bibliometric score (dependent variable) and the external reviewer score (independent variable) stratified by gender, position and year. Linear slopes and accompanying p-values combining data from all years were presented by gender and position. Analyses were performed in R version 3.6.0.

## RESULTS

Between 2014-2017 the KI Career Ladder program attracted 1187 eligible applicants, of which 681 were men and 506 were women. The same applicants may have applied multiple times over the years. Thirty-nine men were awarded a position, resulting in an overall success rate of 5.72%, while 23 women were awarded a position, resulting in an overall success rate of 4.55%. The characteristics of the applicants selected for external peer review are listed in Table 1. Overall, there are no statistically significant differences in external reviewer- or bibliometric scores between men and women. The number of men and women retained at each step, including eligible applicants, those selected for external peer review, those selected for interview, and those finally selected for funding/positions are depicted in Figure 1 and 2.

**Table 1.**
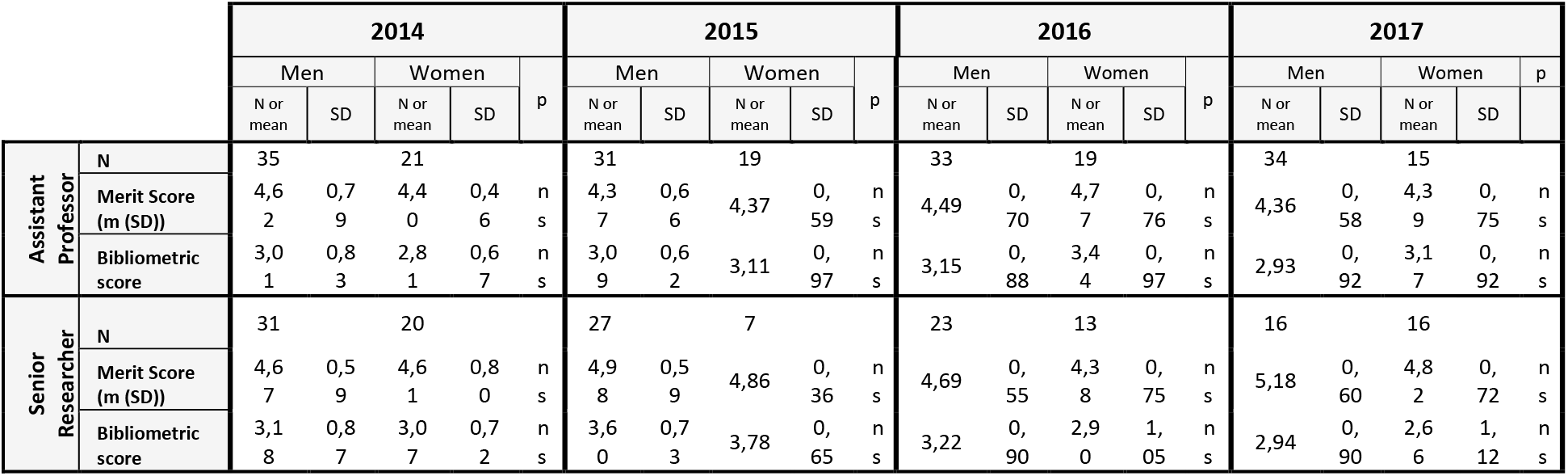
Basic characteristics of the applicants selected for external peer review. No significant differences in bibiometric score or overall merit score between men and women.

**Figure 1.**
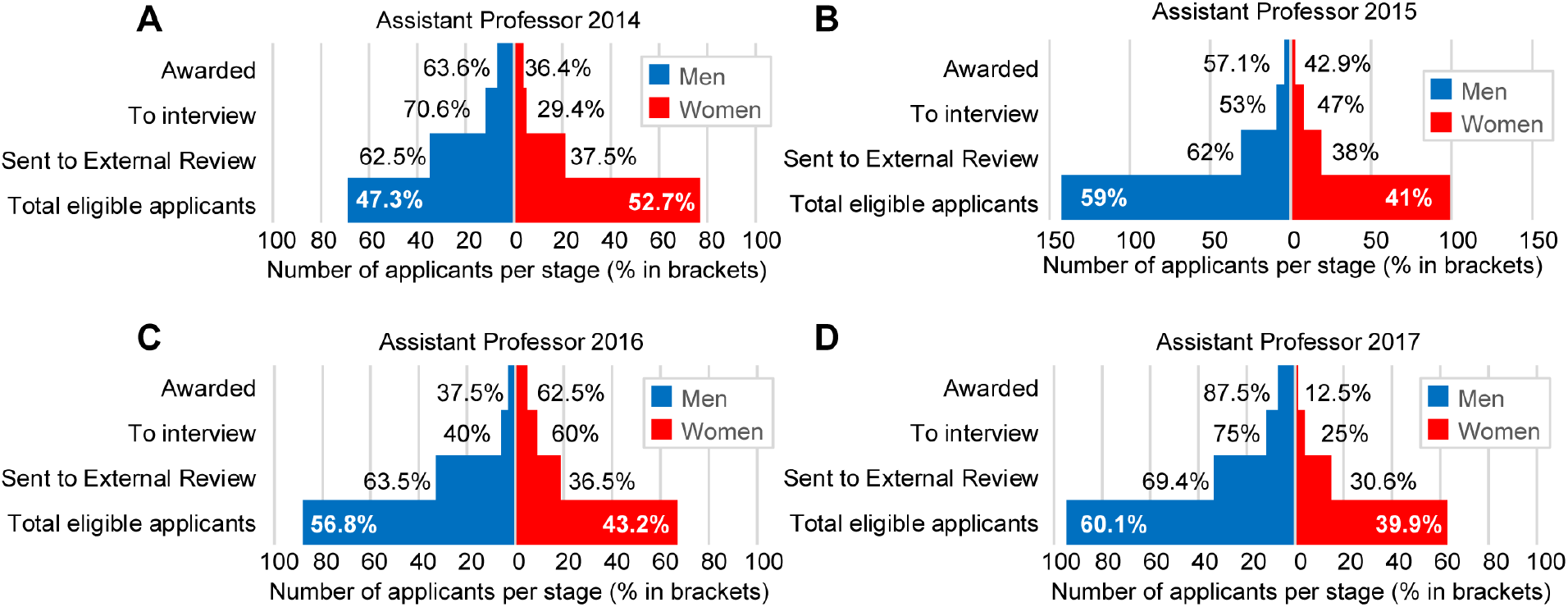
Number of applicants, and proportions, at each step of the recruitment process for the career ladder positions at Karolinska Institutet, divided by year. (A: 2014, B: 2015, C: 2016, D: 2017) for applications to Assistant Professor positions.

**Figure 2.**
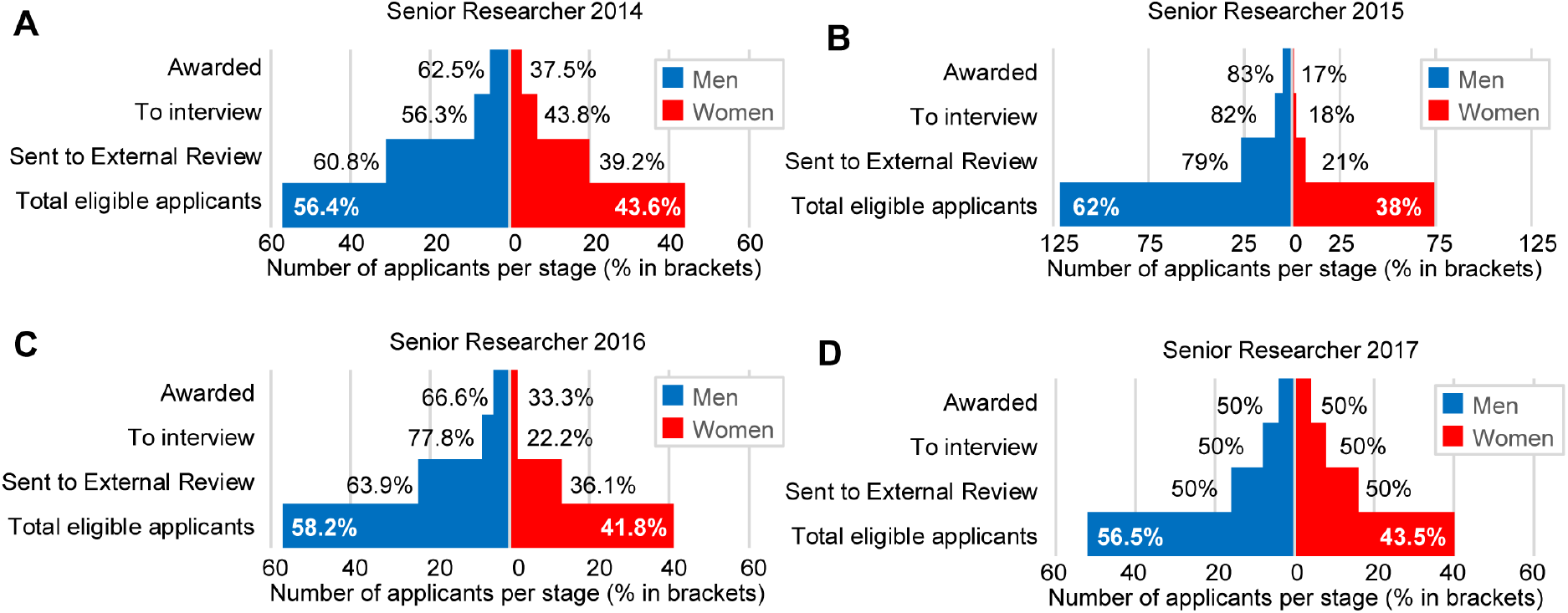
Number of applicants, and proportions, at each step of the recruitment process for the career ladder positions at Karolinska Institutet, divided by year. (A: 2014, B: 2015, C: 2016, D: 2017) for applications to Senior Researcher positions.

### Linear regression results

Positive associations were seen between composite bibliometric scores and external reviewer scores across all years for male applicants (n=132) applying to Assistant Professor positions within the KI career ladder (Figure 3). For women applying to the same positions (n=75), only the last two years (2016-2017) showed a similar positive association (Figure 3 C-D) while the first two years (2014-2015) showed no association at all between composite bibliometric score and external reviewer scores (Figure 3 A-B). This confirms a previous study which analyzed the 2014 Assistant Professor applications using a slightly different composite bibliometric score computation method ^8^.

**Figure 3.**
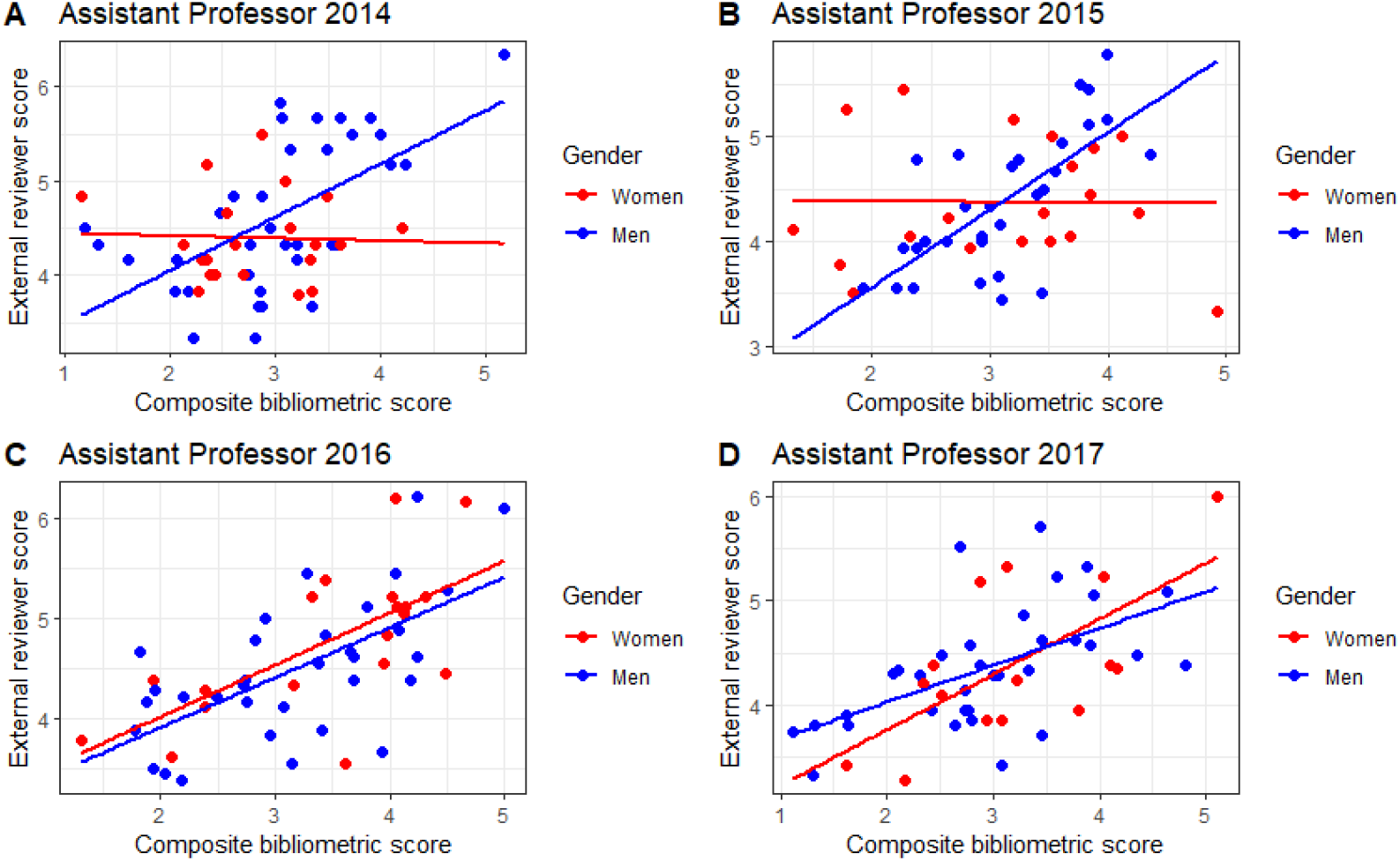
Linear regression associations between composite bibliometric scores and external reviewer scores received on merits, stratified by gender, and divided by year. (A: 2014, B: 2015, C: 2016, D: 2017) for applications to Assistant Professor positions.

For applications to Senior Researcher positions, positive associations were seen for all men (n=96) and women (n=57) across all years (Figure 4).

**Figure 4.**
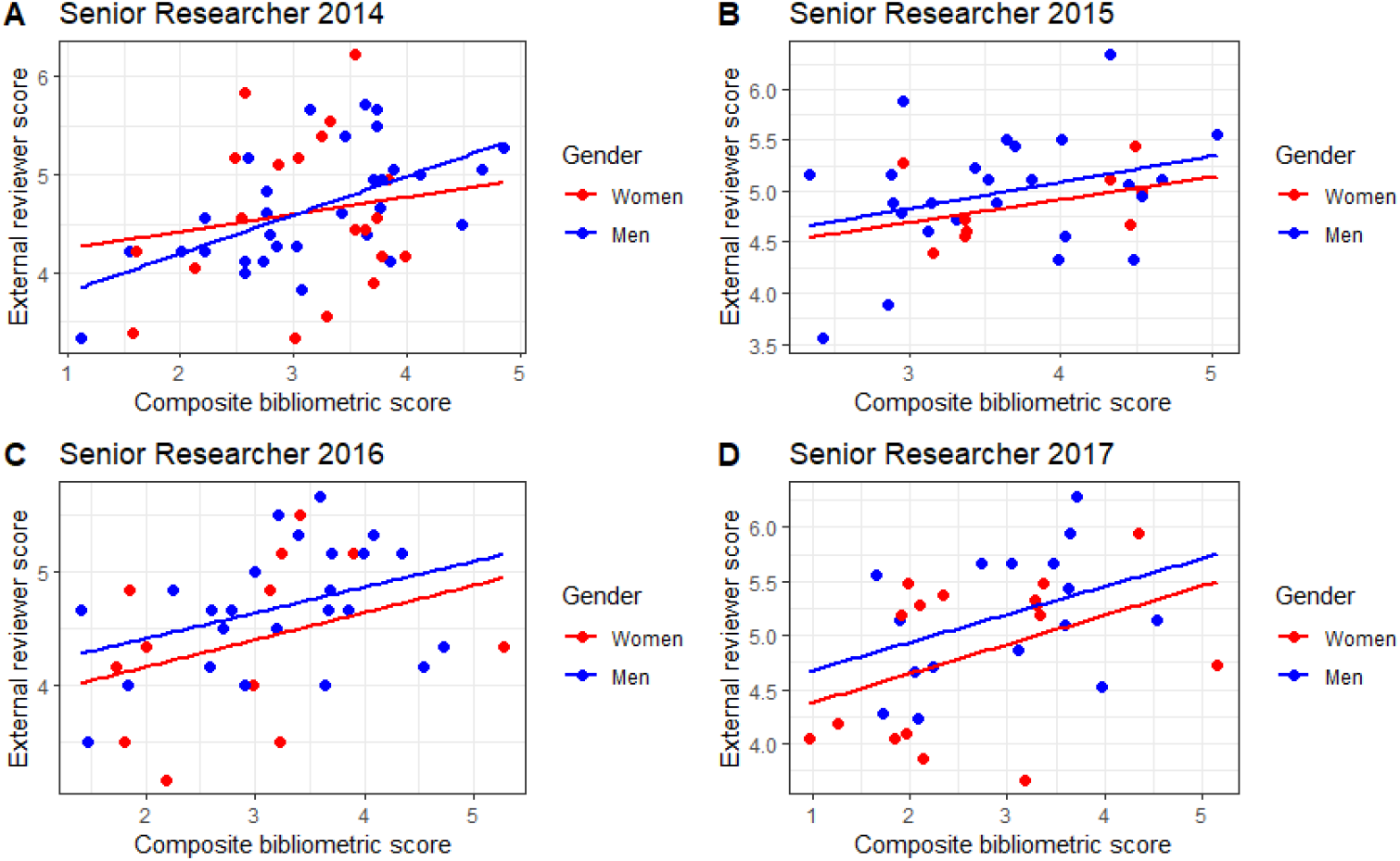
Linear regression associations between composite bibliometric scores and external reviewer scores received on merits, stratified by gender, and divided by year. **(A: 2014, B: 2015, C: 2016, D: 2017)** for applications to Senior Researcher positions.

However, for the last three years (2015-2017) male applicants seemed to have a higher intercept level than women (Figure 4 B-D).

The data for year 2015 deviate from other years, with a sharp increase in the number of applications to both Assistant Professor positions and Senior Researcher positions. This is explained by a change in the eligibility criteria in 2015, now allowing applications from persons holding a permanent position (lab manager, project leader etc.), while these applicants had been excluded (ineligible) in 2014.

Finally, data were pooled for all years for each position, and slopes were compared between men and women (Figure 5). The trend is similar for Assistant Professors (Figure 5A) and Senior Researchers (Figure 5B) in which all slopes had significant positive directions. However, men had consistently steeper slopes and a stronger statistical association between composite bibliometric score and external reviewer score, compared to women (Table 2). In other words, for each additional point in composite bibliometric score, women applying for Assistant Professor positions receive only 58% of the external reviewer score that men receive, and women applying for Senior Researcher positions receive only 79% of the score men receive.

**Table 2.** Linear regression results for associations between composite bibliometric scores and external reviewer scores.

|  | Assistant Professor 2014-2017 |  | Senior Researcher 2014-2017 |  |
| --- | --- | --- | --- | --- |
|  | Slope | P-value | Slope | P-value |
| Men | 0.50 | 8.4E-14 | 0.28 | 7.0E-05 |
| Women | 0.29 | 0.0004 | 0.22 | 0.028 |

**Figure 5.**
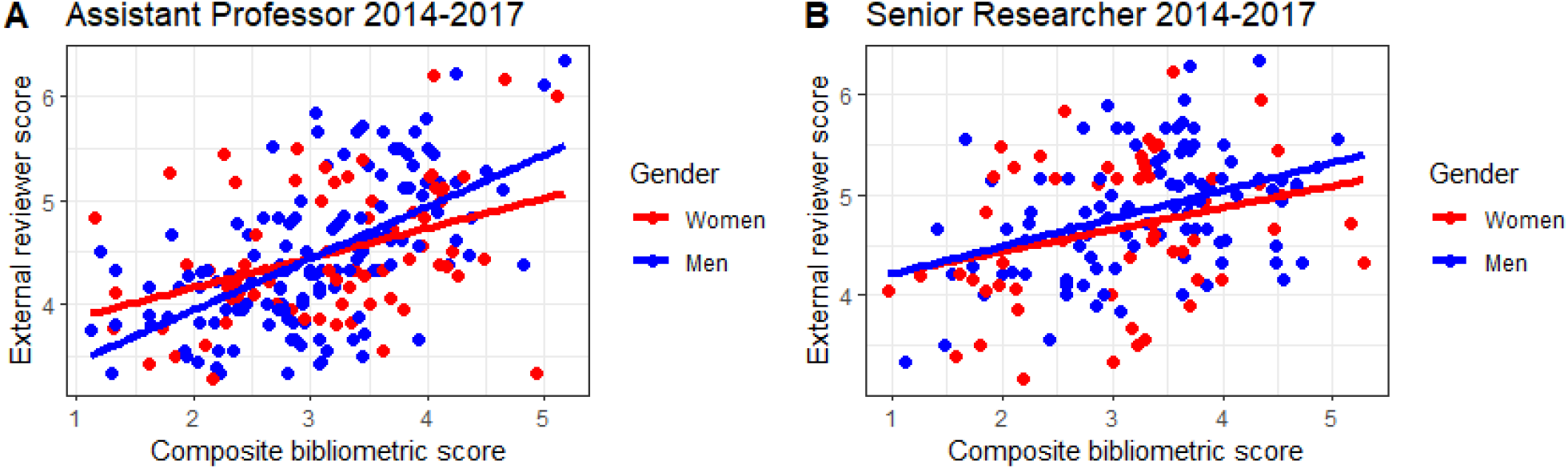
Linear regression associations between composite bibliometric scores and external reviewer scores received on merits combined for all years (2014-2017) and stratified by gender for applications for A: Assistant Professor, and B: Senior Researcher positions.

## DISCUSSION

In this paper, we analyzed gender distributions across a three-step recruitment process for group leader positions at KI (Assistant Professor and Senior Researcher) over a period of four years. We demonstrated clear differences comparing objective quantified productivity and external reviewer scores for merits between men and women, where women on average received lower scores for equal bibliometric achievements. Thus, gender bias in recruitments to higher academic positions is likely to continue to reinforce the unbalanced numbers in professorships in Sweden. When assessing the data for Assistant Professors there is a striking difference in the peer review process comparing 2014-2015 to 2016-2017, because gender differences are apparent in the first years but not in the latter. However, there was no change in the instructions sent out to external reviewers (Appendix) that could explain this deviation. There were no instructions concerning gender bias, nor suggestions as to how to deal with implicit bias. In light of the data presented here, and general awareness of the challenges, KI has started to talk about implicit bias (https://staff.ki.se/assessment-bias-and-career) and is planning to launch a web training program for all reviewers. This effort is a step in the right direction, hopefully with more to come.

Fortunately, there is much data available offering guidance for the construction of a quality-controlled peer-review process. Peer reviewers for NIH applicants show low agreement on the same application ^13^, and a recent preprint in PsyArXiv concluded that at least 12 reviewers per application are needed in order to obtain reliable scores ^14^. KI currently uses six external reviewers. Finland has launched a program in which governmental agencies join forces to establish a standardized template, to be used across all funding bodies, for evaluating researchers in a fair and equal way (https://avointiede.fi/sites/avointiede.fi/files/Vastuullinen-arviointi-luonnos_1.pdf), which may also standardize procedures and remove the impact of bias. Based on the bias we identified here, and previous exhaustive work showing biased review processes ^8–10,15^ we propose a three-step semi-blinded review process (Figure 6). We suggest that project proposals should be assessed blinded to applicant gender, merits could be quantified using the composite bibliometric score, and a reviewer would integrate these two components into an overall assessment. It is important to note that while we consider this to be a possible improvement over current practice, bibliometry itself is biased as well ^11^ and may contribute to continued discrepancies. To address this, the composite bibliometric score could also be corrected by a field-specific “bias factor”.

**Figure 6.**
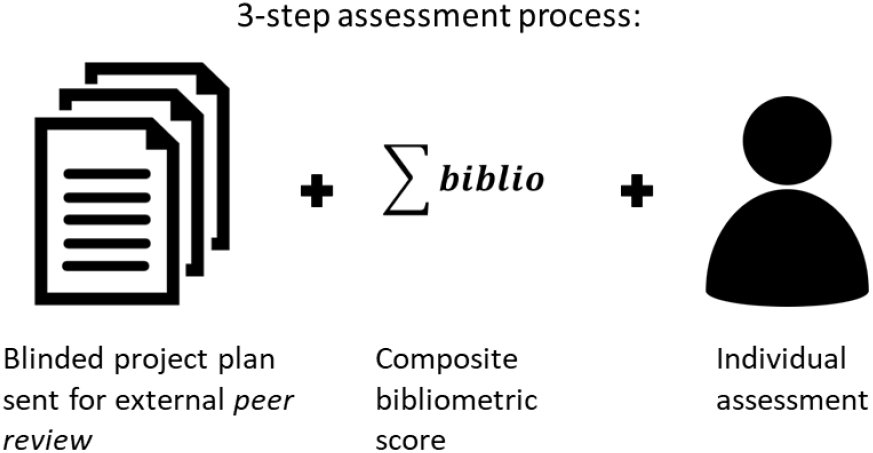
Proposed three-step review process to minimize impact of implicit bias. Project plans can be reviewed blinded by reviewers when applicants are junior, eliminating risk of implicit bias. A composite bibliometric score is calculated to support assessment of past productivity and impact, reducing bias (some metrics have been shown to be unfairly biased by gender). An individual assessment, performed by a peer reviewer, integrates scores from the project plan and the composite bibliometric score, to assess feasibility of the project. Together, these three steps reduce bias while allowing for an assessment of a project’s innovativeness and an applicant’s competence to execute the project.

The difference in success rates for men and women vary widely across position and year, but is higher for men in general. Furthermore, in individual years, the proportion of men or women in the recruitment process is not maintained throughout individual steps, and the proportion of women tends to decrease at each consecutive step (Figure 1 and 2), an expected result if bias is greatest at the top of the competition (Figure 5). If women constitute 53% of the eligible applicant pool, one would expect 53% of the positions to be awarded to women. Local variation is expected, but consistently higher success rates for men throughout the assessment process suggest men are consistently better, which is not supported by our data. Our analysis using the composite score clearly show that this is not the case, and instead men are more highly rewarded for equal merits. There is a fundamental flaw in the “meritocracy” when demographic groups are eliminated as a consequence of not reflecting the norm in academia ^16^.

Sweden is one of the most gender equal countries in Europe ^17^, yet only 25-28% of professors are women ^3,5^. Comparing the percentages of female professors, Sweden is in 13^th^ place, and instead Bulgaria and Latvia lead with 45-54% female professors (“Grade A” positions) ^4,5^. In the 1950’s in Sweden, men and women were equally highly educated, but since the 1960’s more women than men have been highly educated with a difference of 6 percentage points. Currently 39% of women born in 1976 were highly educated at the age of 40 years, while only 23% of men were highly educated. Between 46-49% of doctoral students are women ^3^. In 2017, within medical and health sciences, agricultural and veterinary sciences and social sciences: 59% of doctoral degrees were awarded to women and 41% to men. Thus, education levels cannot explain the dearth of female professors in Swedish universities. Using data from the Swedish Research Council (known as Swedish Medical Research Council at the time), Wennerås and Wold showed in 1997 that female applicants for a postdoctoral fellowship had to be 2.5 times more productive than their male peers to be considered equally merited ^10^. Since then, numerous studies worldwide have demonstrated bias in almost all processes impacting on career progression including hireability of men vs women with identical merits ^7^, desire to mentor men vs women with identical merits ^6^, and probability of citation of similar papers authored by men or women ^6,11^. Over twenty years later, bias against women and non-European men is still significant in the recruitment of junior faculty (Assistant Professors) to Principle Investigator positions at KI ^8^, the results of which we confirm and extend here. Although awareness of bias is increasing, and processes are being implemented to prevent bias from impacting negatively on career progression, recent data show that expectations of brilliance in different academic disciplines still correlate with gender distributions in some fields ^16^. Our data similarly suggest that implicit bias may be more prevalent in the highest tier of competition, slowing the achievement of parity in professorships in academia.

Because citations themselves are biased ^11^, the composite biblimetric score is not a completely unbiased assessment of productivity, and should be used with caution. Further, the suitability of assessing individuals based on publication metrics is a heavily debated issue. However, considering the challenges in fairly assessing women, this is an improved measure of productivity compared to reviewer assessment, which we show is biased.

To conclude, in order to attract and maintain the best scientists in the academic career track, all individuals should have equal opportunities and be reviewed in transparent systems on equal terms. Ensuring a meritocracy with the best individuals in academia will require more work to ensure quality-controlled recruitment and assessment procedures.

## ACKNOWLEDGEMENTS

We thank Karolinska Institutet for supporting this work financially and with data, as well as for extensive discussions of how to improve the future recruitment processes to prevent bias. We are grateful for support from the KI Junior Faculty Steering Group, the KI Leadership, funding from KI Junior Faculty (awarded by the KI Board of Research to Junior Faculty). We also thank the KI Library for assistance in analysis of bibliometric data.

## Contributions

ERA: Conceptualization, Data Curation, Formal Analysis, Investigation, Methodology, Project Administration, Visualization, Writing – Original Draft Preparation, Writing – Review & Editing

CH: Writing – Review & Editing

SH: Conceptualization, Data Curation, Formal Analysis, Investigation, Methodology, Project Administration, Visualization, Writing – Original Draft Preparation, Writing – Review & Editing

